# Myo1e knockout in the adult podocytes leads to proteinuria but has less severe consequences for kidney function than Myo1e loss during renal development

**DOI:** 10.1101/2022.03.03.482915

**Authors:** Sharon E. Chase, Mira Krendel

## Abstract

Myosin 1e (Myo1e) is expressed in specialized epithelial cells in the kidney (podocytes) and plays an important role in renal filtration. Knockout of Myo1e in mice and mutations in the *MYO1E* gene in humans cause proteinuria and disrupt the ultrastructure of glomeruli, the initial segments of renal nephrons that are responsible for selective excretion of waste products without the loss of proteins from the bloodstream. Previous studies have demonstrated that the loss of Myo1e early in development results in severe defects in renal function in mice and may lead to end-stage renal disease in patients homozygous for *MYO1E* mutations. However, little is known about the effects of Myo1e loss later in life. In this study, we used inducible knockout of Myo1e in mouse podocytes to examine the effects of Myo1e loss from the adult kidneys. We have found that Myo1e loss after the completion of renal development causes proteinuria and podocyte defects but the effects are milder and more variable compared to the effects of Myo1e knockout in developing podocytes. These findings indicate that Myo1e plays an important role in podocyte development and differentiation but may also contribute to the maintenance of the glomerular barrier in the adult kidneys.

## Introduction

The first step in the filtration of waste products by the kidney is the passage of fluid and solutes through the renal glomerulus, a collection of fenestrated (porous) capillaries covered with specialized epithelial cells called podocytes. The glomerular filtration barrier allows water and dissolved small molecules to pass into the urinary space while selectively retaining blood cells and proteins in the bloodstream. Glomerular epithelial cells (podocytes) represent one of the main components of the selective filtration barrier. Podocyte loss or dysfunction leads to the disruption of selective filtration and result in proteinuria, which may lead to end stage kidney disease if left untreated. Podocytes form actin-rich processes (foot processes) that cover the entire surface of glomerular capillaries, and the integrity of the podocyte actin cytoskeleton is crucial for normal glomerular filtration (Faul et al., 2007).

Myosin 1e (Myo1e) is a member of the myosin family of actin-dependent molecular motors (Bement et al., 1994; Krendel and Mooseker, 2005) that is expressed in podocytes (Krendel et al., 2009). Myo1e knockout mice exhibit proteinuria starting at 3 weeks after birth (Krendel et al., 2009). Mutations in the *MYO1E* gene have been linked to pediatric familial focal segmental glomerulosclerosis (FSGS) in humans, further highlighting the role of Myo1e in glomerular functions (Mele et al., 2011; Sanna-Cherchi et al., 2011). Both Myo1e-null mice and patients with the MYO1E mutations exhibit podocyte foot process effacement (flattening) and characteristic changes in the GBM (thickening and splitting). Similar defects are observed in mice with the podocyte-specific knockout of Myo1e (Chase et al., 2012), indicating that podocytes represent the primary target of the Myo1e activity in the kidney. Since patients homozygous for Myo1e mutations develop FSGS at an early age, it is possible that Myo1e activity plays important roles during glomerular development, for example, during formation of podocyte foot processes and slit diaphragms.

In addition to its role in glomerular filtration, Myo1e contributes to progression of breast cancer and may influence tumor metastasis (Ouderkirk and Krendel, 2014). In order to determine whether Myo1e may be used as a target for developing metastasis-targeting therapeutics, it is important to know whether inhibition of Myo1e functions in adult patients is likely to have detrimental consequences on renal functions. To this end, we set out to determine whether Myo1e expression in podocytes is required to maintain the intact filtration barrier in the adult kidney. We have created a mouse model in which Myo1e knockout in podocytes can be induced by administration of doxycycline. This mouse model contains two Myo1e alleles. One is a constitutive knockout allele (Myo1e^-^) and the other is a conditional knockout allele (Myo1e^Flox^), which contains two loxP sites. The loxP sites allow the excision of the Myo1e exon 4 by Cre recombinase and disrupt Myo1e expression by the same mechanism that was originally used to create Myo1e^-/-^ mice, namely, by creating a stop codon near the N-terminus of the protein(Krendel et al., 2009). Thus, under the control conditions, the mice used in this study are heterozygous for Myo1e and have normal kidney functions, as previously described for Myo1e^+/-^ animals (Krendel et al., 2009). The mouse model used in this study also contains two transgenes, Podo-rtTA and tetO-Cre, which encode the reverse tetracycline transactivator and Cre recombinase, respectively (Perl et al., 2002; Shigehara et al., 2003). The Cre recombinase expression is under the control of a tetracycline-responsive promoter, which requires the binding of rtTA complexed with doxycycline to induce protein expression. The rtTA protein is expressed under the control of the podocin promoter, limiting its expression to podocytes. Doxycycline administration results in Cre expression in podocytes, generating mice with a podocyte-targeted knockout of Myo1e (Fig.1).

**Figure 1.**
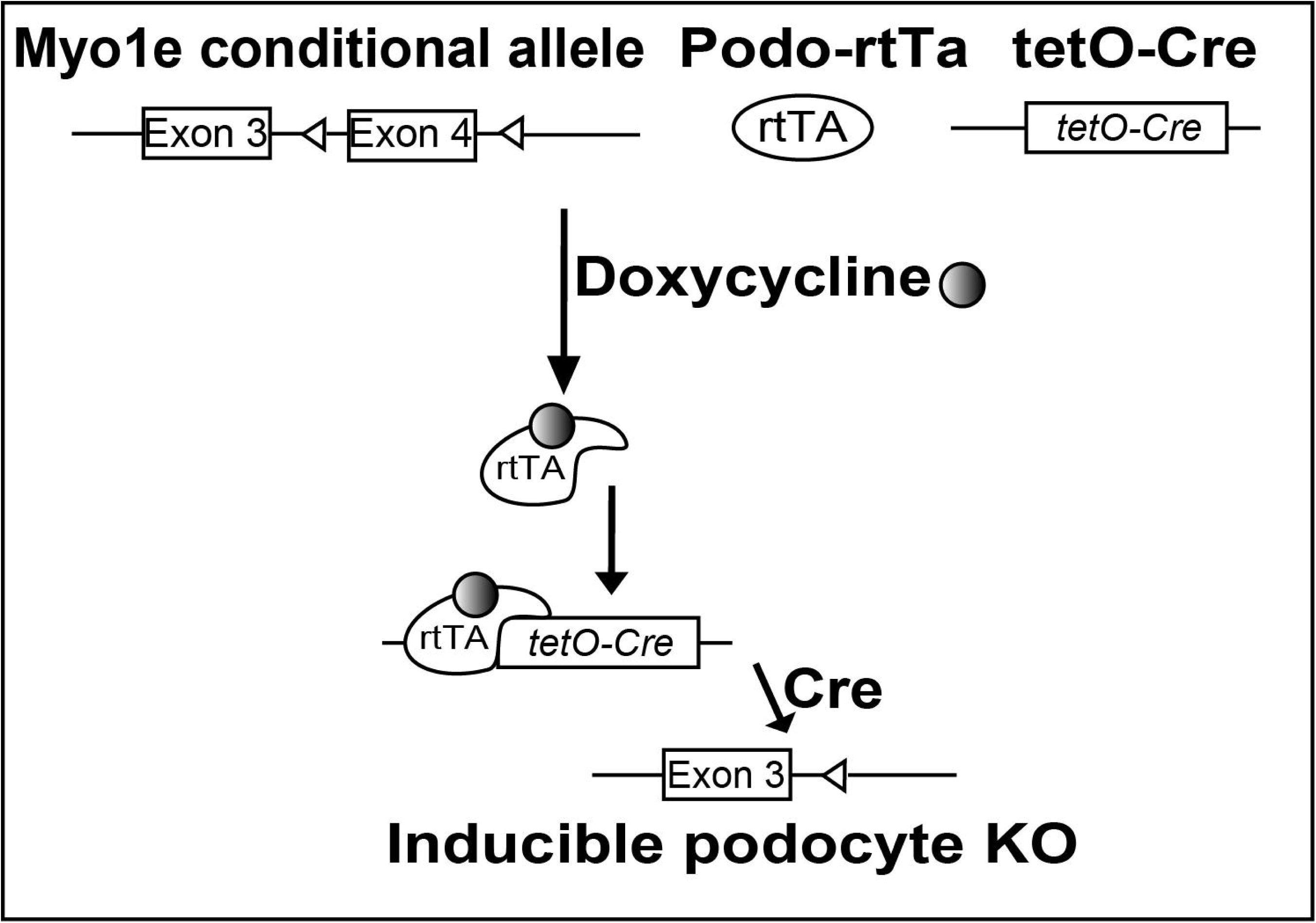
Schematic of the doxycycline-inducible knockout of Myo1e in podocytes. Myo1e KO is accomplished by Cre recombinase-mediated removal of Exon 4 in the mouse *Myo1e* gene, which results in premature termination of translation. Cre recombinase expression off of the doxycycline-inducible *tetO* promoter requires expression of doxycycline transactivator, rtTA, which is expressed from the Podo-rtTA transgene, under the control of the *NPHS2* (podocyte-specific) promoter.

Using doxycycline treatment, we induced disruption of Myo1e expression in podocytes either during prenatal development or in the adult. We confirmed Myo1e loss using immunostaining, measured protein excretion in urine, and examined glomerular ultrastructural changes following Myo1e knockout. We have also tested how Myo1e loss in the adult affects susceptibility to acute and chronic renal disease using low dose LPS injection as a model of transient injury and doxorubicin injection as a model of chronic nephropathy. Our findings indicate that Myo1e loss from the adult kidneys disrupts glomerular integrity; however, the effects of Myo1e knockout in adult podocytes on kidney function are highly variable and produce a milder phenotype than the podocyte-targeted perinatal knockout. Overall, our findings indicate that Myo1e inhibition in the adults for therapeutic purposes may be tolerated if accompanied by careful monitoring of renal function.

## Materials and methods

### Animals

Myo1e^-/-^and Myo1e ^Flox/Flox^ mice have been previously described (Chase et al., 2012). These mice were maintained on C57/Bl6 background in our lab. B6.Cg-Tg(tetO-cre)1Jaw/J mice (stock 006234), FVB/N-Tg(NPHS2-rtTA2*M2)1Jbk/J mice (stock 008202), and mTmG mice, B6.129(Cg)-*Gt(ROSA)26Sor*^*tm4(ACTB-tdTomato,-EGFP)Luo*^/J, (stock 007676) were purchased from Jackson Labs. All protocols have been approved by SUNY Upstate Medical University’s IACUC under protocol 364. Doxycycline was administered at 2 mg/ml in water with 3% sucrose or in food (625 ppm, Test Diet, Purina Mills). For perinatal treatment, doxycycline was administered to pregnant females beginning at day 13 with continued administration to lactating females until weaning.

*Genotyping* for Myo1e Flox/wt/KO and for tetO-Cre transgene was performed as previously described (Chase et al., 2012). Specifically, primers Myo1e A (TCATGTGTAGCCCAAGCTCACC), Myo1e B (TTCCGCTTACGGTGGAAATG), and Myo1e C (ACTCATTCTGTCATCTGACTCCACC), were used for Myo1e genotyping, and primers Cre For (GCATAACCAGTGAAACAGCATTGCTG) and Cre Rev (GGACATGTTCAGGGATCGCCAGGCG) were used to detect tetO-Cre transgene. Primers oIMR8111(GAACAACGCCAAGTCATTCCGTC) and oIMR8112 (TACGCAGCCCAGTGTAAAGTGGTT) were used to detect rtTA transgene and primers oIMR7338 (CTAGGCCACAGAATTGAAAGATCT) and oIMR7339 (GTAGGTGGAAATTCTAGCATCATCC) were used to detect an internal positive control band according to the Jackson Labs protocol for rtTA genotyping. Primers mTmG#1 (CTCTGCTGCCTCCTGGCTTCT), mTmG#2 (CGAGGCGGATCACAAGCAATA), mTmG#3 (TCAATGGGCGGGGGTCGTT) were used to genotype mTmG mice (the presence of a 330 bp band indicates that mTmG allele is not present, the presence of a 250 bp band designates the presence of the mTmG transgene).

Urinary albumin measurements were performed using Coomassie-stained SDS-PAGE with BSA standards as previously described (Chase et al., 2012). Urinary creatinine was measured using the Creatinine Companion kit, Ethos Biosciences, Philadelphia, PA, cat#1012.

### Immunohistochemistry and histology

For immunofluorescence staining, fresh (non-perfused) kidneys were frozen in OCT for cryosectioning. Cryosections were fixed with 3% paraformaldehyde/PBS (15 min.), permeabilized with 0.25% Triton X-100/PBS (3 min.), and stained using the following antibodies: anti-synaptopodin (clone G1D4, Genetex), anti-myo1e (Skowron et al., 1998). For histological staining, formalin-fixed, paraffin-embedded kidneys were sections and stained with Harris Hematoxylin and Eosin (Luna et al., 1968), Alcian Blue-PAS (University of Rochester Medical Center, 1979), or Masson’s Trichrome (Luna et al., 1968). Images of immunofluorescently labeled kidney sections were collected using a Perkin-Elmer UltraView VoX Spinning Disc Confocal system mounted on a Nikon Eclipse Ti microscope. Images of histology slides were collected using a Canon EOS Rebel DS126311 camera on an Olympus CH student microscope.

Kidney fixation and processing for electron microscopy were performed as previously described (Krendel et al., 2009). Morphometry on kidney sections was performed using ImageJ software. Five to fifteen micrographs per animal (obtained from different glomeruli) were included in the analysis, and 1-2 animals were analyzed for each genotype. Glomerular basement membrane (GBM) in each electron micrograph was manually outlined in ImageJ to determine its area. The total area of all regions of the basement membrane present in each micrograph was divided by the total length of the basement membrane (measured by drawing a line through the center of each basement membrane segment) to determine its average thickness. Foot process number was manually counted and divided by the length of the basement membrane.

### Glomerular isolation and imaging

Glomeruli were isolated from mTmG mouse kidneys using a sieving procedure. Briefly, kidneys were minced in cold PBS and ground through 180 um, 100 um, and 71 um sieves, using cold PBS to rinse the sieves. Glomeruli were collected from the top surface of the 71 um sieve and washed in cold PBS by centrifugation at 500g for 5 min., fixed using 4% PFA in PBS, washed again in cold PBS, placed in glass-bottom 35 mm dishes (Mattek, Ashland, MA), and imaged using the Perkin-Elmer UltraView VoX Spinning Disc Confocal system mounted on a Nikon Eclipse Ti microscope.

### Adriamycin nephropathy model

3-4 weeks after the end of the administration of doxycycline, Adriamycin (doxorubicin) was administered via retroorbital injection under anesthesia at 15 ug/g of mouse weight. Urine samples were collected weekly for 6 weeks after the injection of doxorubicin.

### LPS injury model

3 weeks after the end of the administration of doxycycline, LPS (Invivogen, Ultrapure) was administered by intraperitoneal injection at 10 ug/g of mouse weight at a concentration of 0.5 mg/ml in sterile saline. Urine samples were collected prior to the injection of LPS and at days 1, 2, 3, 4, 5, and 7 post-injection.

## Results

### Inducible knockout of Myo1e during renal development causes proteinuria and defects in glomerular organization

We have previously demonstrated that mice with the podocyte-specific knockout of Myo1e (using NPHS2 promoter-driven Cre recombinase transgene, also known as PodoCre) develop moderate proteinuria, podocyte effacement, and GBM defects (Chase et al., 2012). In order to verify that the inducible knockout of Myo1e using the doxycycline system resulted in the outcomes consistent with our previous observations, we first used doxycycline treatment to induce Myo1e knockout during prenatal and postnatal development. Pregnant females were given doxycycline in drinking water, and doxycycline administration was continued until weaning. Following weaning, the pups were genotyped, and mice that contained a complete set of transgenes required for the Myo1e knockout (Myo1e^-/Flox^ tetO-Cre NPHS2-rtTA) were designated as “experimental” while their littermates that have also received doxycycline but were missing the rtTA or tetO-Cre transgenes were used as controls. Immunostaining for Myo1e showed that experimental animals exhibited loss of Myo1e in the glomeruli (Fig. 2A), although the extent of Myo1e loss was variable among individual glomeruli.

**Figure 2.**
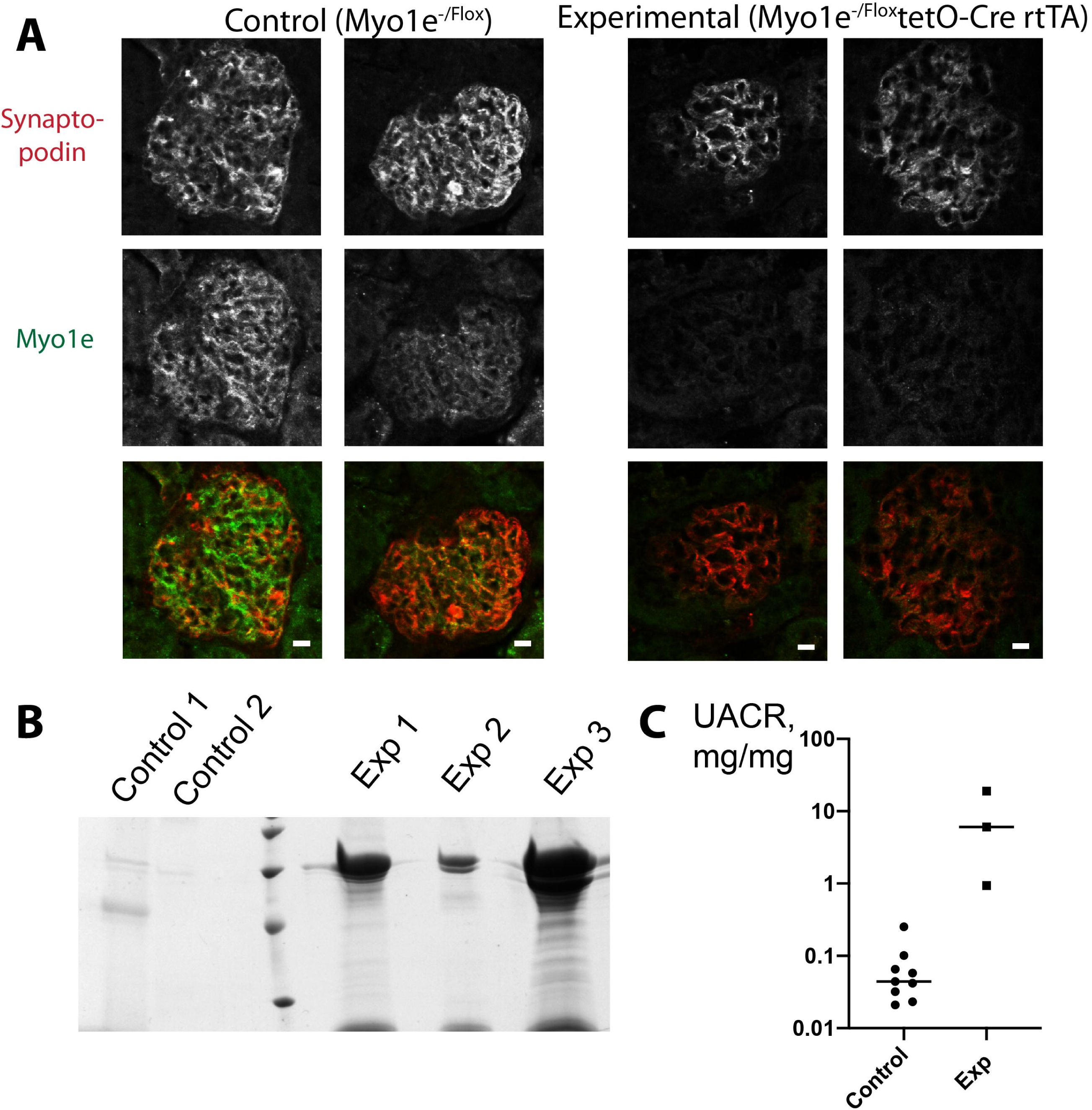
Perinatal knockout of Myo1e using the doxycycline-inducible system results in proteinuria. Control and experimental mice (carrying one KO and one floxed allele of Myo1e together with the rtTA and tetO-Cre transgenes) were treated with doxycycline perinatally by providing doxycycline to females during pregnancy and lactation periods. A, Immunohistochemical labeling of Myo1e (green) and podocyte marker synaptopodin (red) in kidney cryosections of mouse kidneys from 7-week-old mice. In the control mouse kidneys, myo1e colocalizes with synaptopodin in podocytes. In the KO (experimental) kidneys, myo1e staining is no longer observed in podocytes. Scale bar, 10 um. B. Representative urine samples from 5–7-week-old control and myo1e-KO mice (5 ul) were separated by SDS-PAGE and stained with Coomassie Blue. Control 1 = Myo1e^-/Flox^ tetO-Cre, control 2 = Myo1e^-/Flox^. C, Urinary albumin/creatinine ratio is elevated in urine samples from 5–7-week-old myo1e-KO mice compared to the control mice.

Analysis of urine samples from the control and experimental animals showed that experimental mice developed moderate proteinuria by the age of 6-7 weeks (Fig. 2B,C). The extent of proteinuria as measured by the urinary albumin/creatinine ratio was similar to that previously observed in the Myo1e^-/Flox^ PodoCre mice that we had used to obtain targeted knockout of Myo1e in podocytes (Chase et al., 2012). Similarly to our previous observations in the podocyte-targeted Myo1e knockout model, the experimental mice showed variable response to the treatment, with some animals maintaining normal protein filtration (Fig. 2C). This likely reflects the variability in the Cre recombinase expression efficiency and the extent of Myo1e loss.

We used transmission electron microscopy to examine glomerular organization in the control and experimental mice. The experimental mice that showed detectable proteinuria were selected for electron microscopy analysis. At the ultrastructural level, their kidneys were characterized by podocyte foot process effacement and localized widening of the GBM (Fig. 3). The GBM exhibited changes typical of Myo1e-knockout animals and patients with *MYO1E* mutations, namely, the presence of electron-lucent areas and widening and bulging out of the GBM.

**Figure 3.**
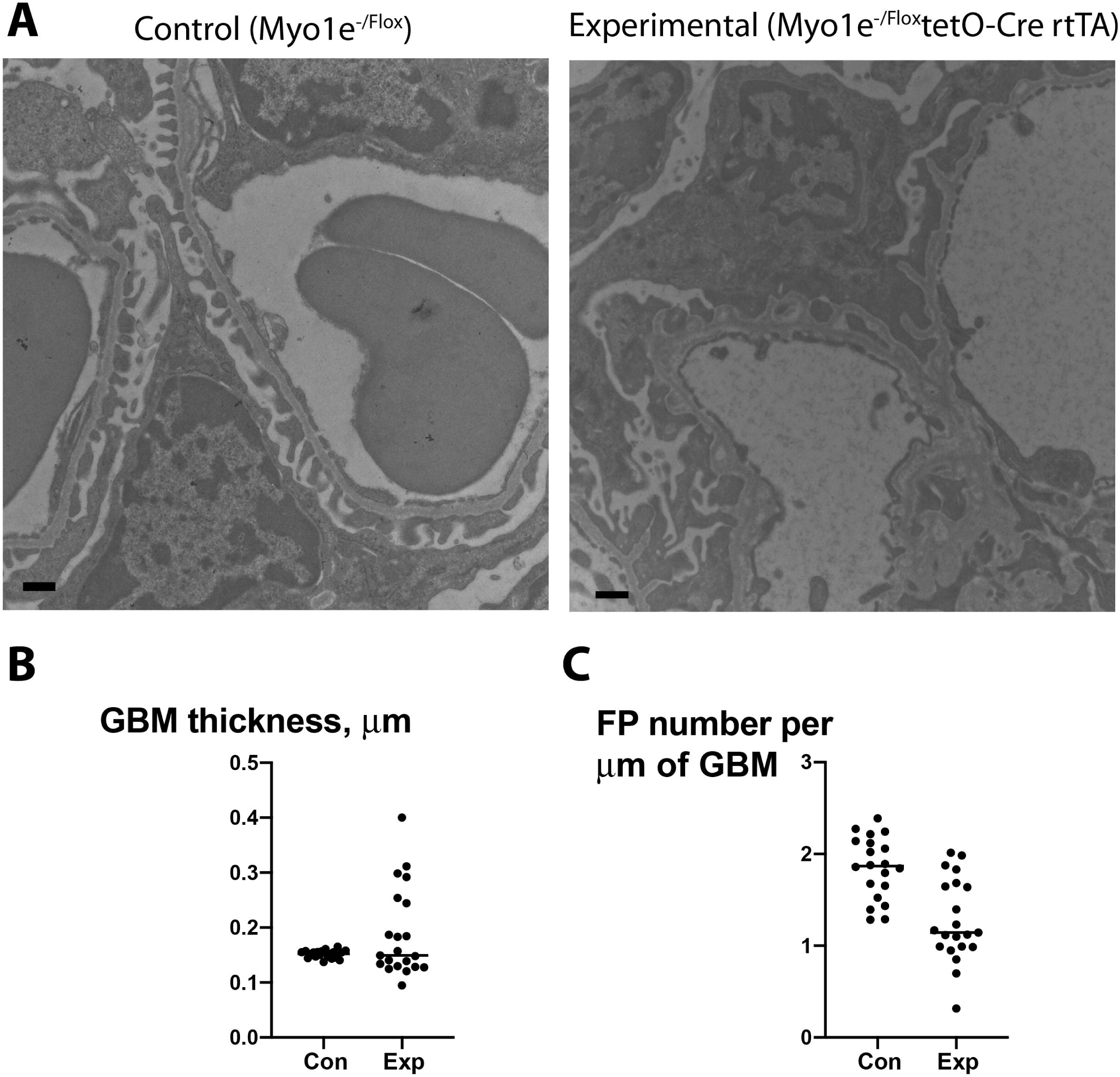
Knockout of Myo1e during renal development results is defects in glomerular ultrastructure. A. Electron micrographs of kidney samples from control and experimental mice treated with doxycycline during the perinatal period demonstrate that glomeruli of Myo1e-KO mice are characterized by foot process effacement and uneven thickening and lamination of the GBM. Scale bars, 2 um. B, C. Quantification of the GBM width and foot process number shows that the GBM width is increased in the Myo1e-KO mice (B, P=0.0465) while foot process density is decreased in the Myo1e-KO mice compared to controls (C, P<0.0001). Each data point corresponds to a single field of view. 1 control and 2 KO mice were used for quantification.

### Induction of the Myo1e knockout in the adult kidney leads to the disruption of selective protein filtration and defects in glomerular ultrastructure

After confirming that the induction of Myo1e knockout in developing glomeruli had effects similar to those previously observed for the constitutive podocyte-targeted knockout, we proceeded to test the effects of Myo1e loss on the established filtration barrier. Experimental mice (Myo1e^-/Flox^ rtTA tetO-Cre) and control mice (Myo1e^-/Flox^ or Myo1e^-/Flox^ tetO-Cre littermates of the experimental mice) were given doxycycline in drinking water for 2 weeks, starting at the age of 8-10 weeks, to induce Cre expression and Myo1e loss from podocytes. Urinary albumin and creatinine were monitored weekly before and after doxycycline treatment, and mice were sacrificed several weeks after treatment to collect kidneys for histological and ultrastructural characterization. In addition, to confirm the efficiency of the doxycycline-induced Cre-mediated recombination, we used mice containing a transgene that encodes a membrane-targeted red fluorescent protein tdTomato and a membrane-targeted green fluorescent protein (mTmG mice). Cells containing this transgene express tdTomato unless subjected to Cre-mediated recombination, which results in a switch to GFP expression (Muzumdar et al., 2007). When mice carrying the mTmG, rtTA, and tetO-Cre transgenes were treated with doxycycline, we observed expression of GFP in glomeruli, with a pattern consistent with podocyte localization (Fig. S1). Doxycycline treatment had no effect on the albumin/creatinine ratio in the mTmG mice.

We found that doxycycline treatment had no effect on myo1e staining, albumin excretion, or albumin/creatinine ratio in control mice (Myo1e^-/Flox^ or Myo1e^-/Flox^ tetO-Cre mice) (Fig. 4). In contrast, immunohistochemical analysis of kidney sections from Myo1e-KO mice (Myo1e^-/Flox^ tetO-Cre NPHS2-rtTA) showed that expression of Myo1e in glomeruli was reduced (Fig. 4A). Loss of the linear Myo1e staining in podocyte foot processes located along the glomerular basement membrane was especially noticeable (see yellow and red arrows in Fig. 4A, indicating colocalization of Myo1e staining with synaptopodin staining in control and lack of colocalization in experimental kidney sections).

**Figure 4.**
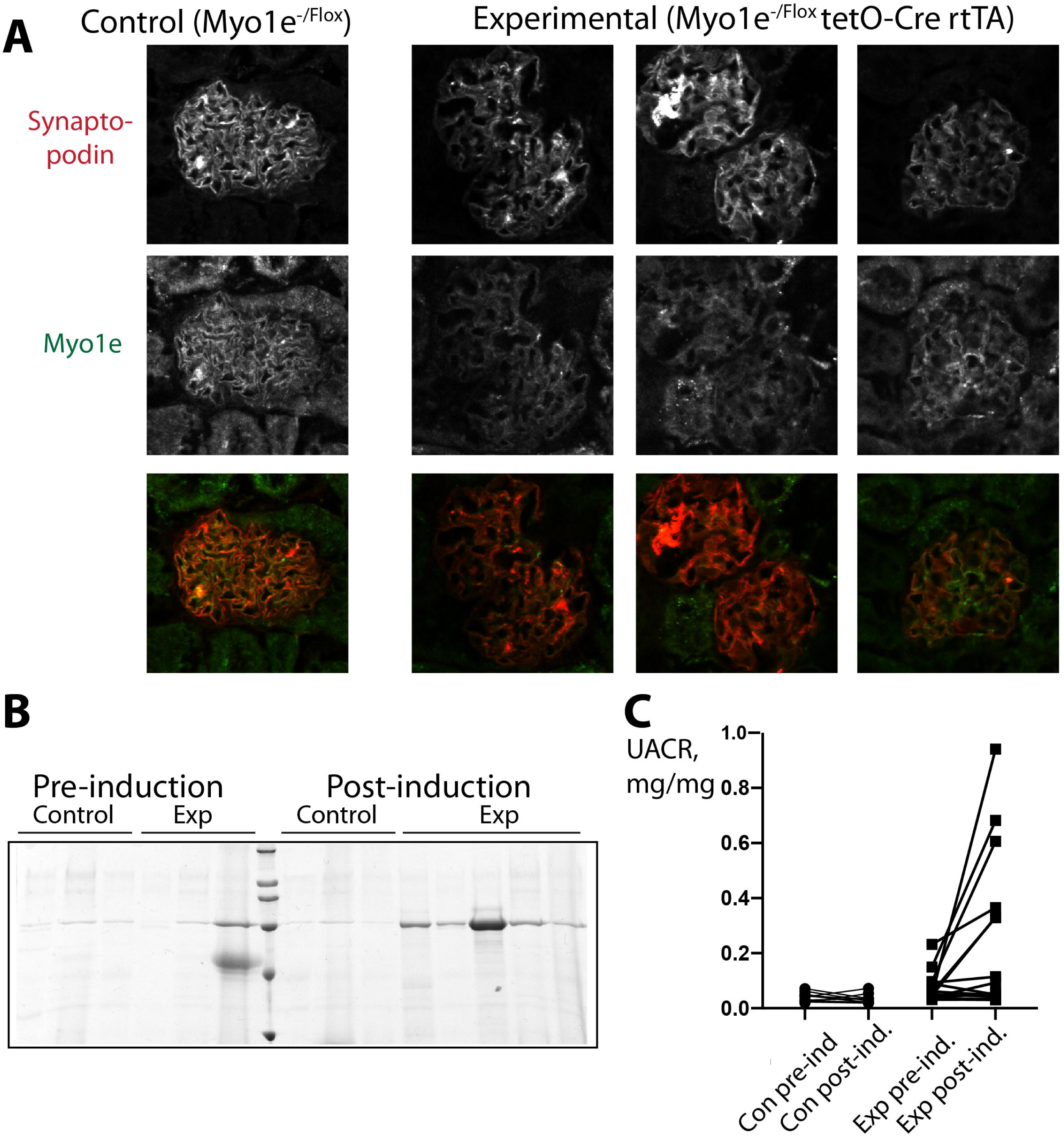
Doxycycline-induced knockout of Myo1e in podocytes of adult mice results in variable extent of proteinuria. Control and experimental mice were treated with doxycycline for 2 weeks starting at the age of 8 weeks. A, Immunohistochemical labeling of Myo1e (green) and podocyte marker synaptopodin (red) in cryosections of kidneys from 21 week-old mice, 10 weeks after the administration of doxycycline. In the control mouse kidneys, myo1e colocalizes with synaptopodin in podocytes. In the KO (experimental) kidneys, myo1e staining is no longer observed in podocytes. Scale bar, 10 um. B. Representative urine samples (5 ul each) from control and myo1e-KO mice (prior to doxycycline treatment and 2 weeks after the end of doxycycline treatment) were separated by SDS-PAGE and stained with Coomassie Blue. C, Urinary albumin/creatinine ratio measured in urine samples from myo1e-KO mice and control mice prior to doxycycline treatment and 5 weeks after the end of doxycycline treatment.

Doxycycline treatment resulted in an increase in urinary albumin excretion in some of the experimental animals (Fig. 4B,C). An increase in albumin excretion was not observed in all experimental animals, likely reflecting the limited efficiency of the Cre-mediated recombination in this system. Indeed, when we examined the efficiency of recombination using mTmG reporter mice, only a fraction of all podocytes exhibited GFP expression, indicative of successful recombination (Fig. S1). The extent of proteinuria in mice where KO was induced after the completion of renal development was less than in mice with the perinatal KO of Myo1e (compare Figures 2 and 4).

When Myo1e KO mice exhibiting proteinuria were subjected to further analysis of glomerular ultrastructure using EM, we found that glomeruli of Myo1e-KO mice were characterized by GBM thickening and foot process effacement (Fig. 5) compared to the control mice. GBM changes in the experimental animals were reminiscent of those previously observed in the mice with constitutive and perinatal/podocyte-targeted Myo1e KO, including bulging areas and electron-lucent areas with basket-weave-like structure (Chase et al., 2012; Krendel et al., 2009; Randles et al., 2016), see also Fig. 3. Upon histological examination, Myo1e-KO mice, even those that exhibited proteinuria, did not demonstrate significant glomerular scarring or interstitial fibrosis (Fig. S2), indicating the lack of wide-spread renal damage. Surprisingly, even the mice that exhibited dramatic changes in the foot process width and the GBM architecture displayed only mild proteinuria. Thus, overall, the loss of Myo1e from adult podocytes does not have the same disruptive effect on renal filtration as the perinatal KO of Myo1e, indicating that during kidney development Myo1e is likely to play an important function in podocyte differentiation that is no longer required once mature podocytes are formed.

**Figure 5.**
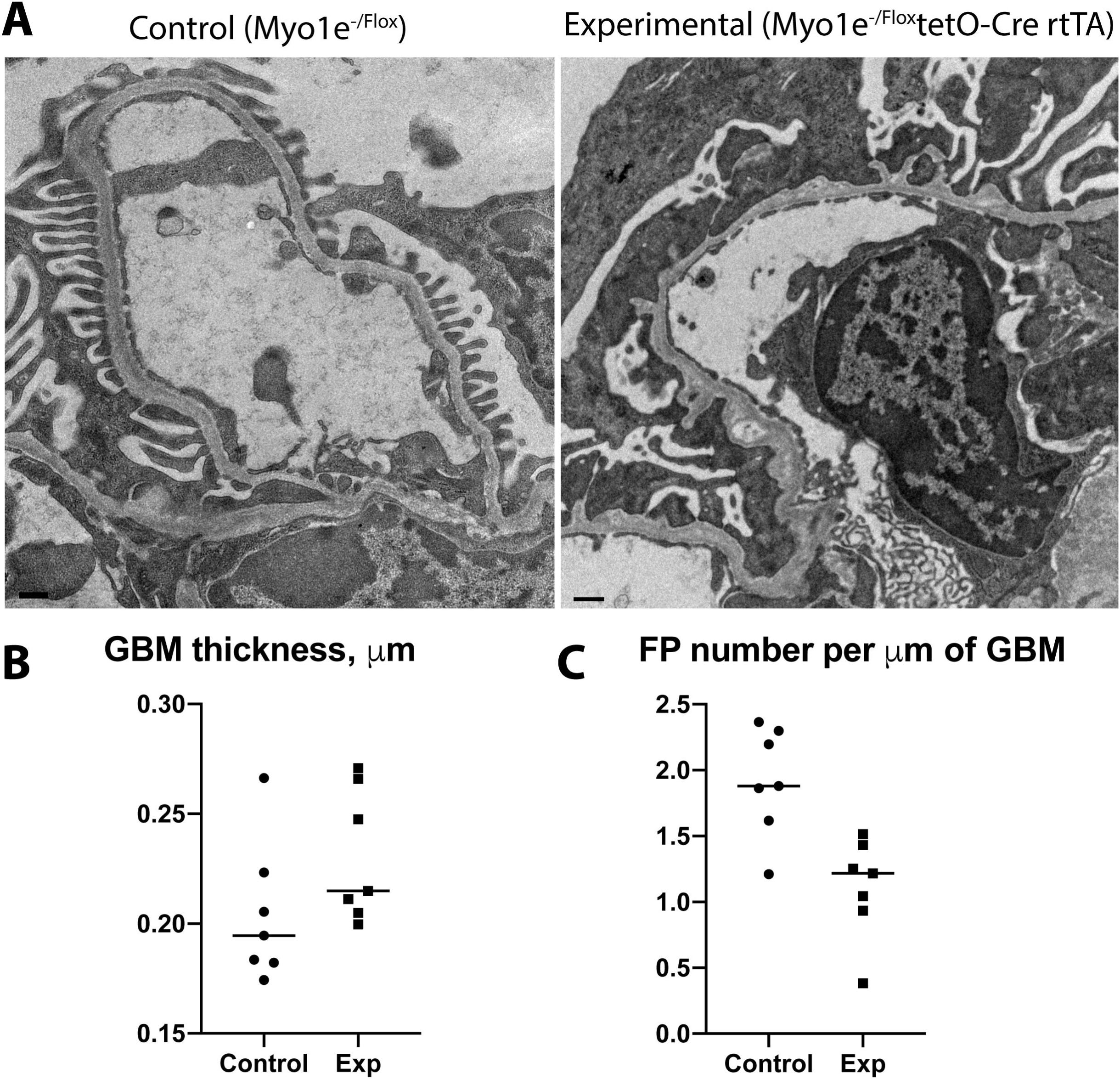
Knockout of Myo1e in adult podocytes leads to disruption of GBM organization and foot process effacement. A. Electron micrographs of kidney samples from Myo1e control and KO mice treated with doxycycline for 2 weeks starting at the age of 8 weeks. Samples were collected from 21 week-old mice. Myo1e-KO mice exhibit foot process effacement and disorganized GBM. B. Measurements of the GBM width from electron micrographs show variability in the GBM width in the Myo1e-KO mice but the differences between control and KO mice are not statistically significant. C. Foot process number is reduced in the Myo1e-KO mice (P=0.0025). Each data point in B and C corresponds to a single field of view. 2 control and 2 KO mice were used for quantification.

### Myo1e knockout in the adult kidney and susceptibility to the LPS-induced proteinuria and Adriamycin nephropathy

We hypothesized that the developmental contribution of Myo1e to shaping podocyte architecture may be recapitulated during the recovery from a transient or long-term kidney injury. To test whether Myo1e loss makes kidneys more susceptible to transient injury or delays recovery, we subjected control and Myo1e-KO mice to a low-dose LPS treatment (Faul et al., 2008). This treatment induces transient proteinuria followed by a rapid recovery. Mice with the induced KO of Myo1e demonstrated a more pronounced increase in proteinuria following LPS administration and slightly elevated proteinuria during recovery post-LPS injection, however, the differences in proteinuria between the control and experimental mice were not statistically significant (Fig. 6).

**Figure 6.**
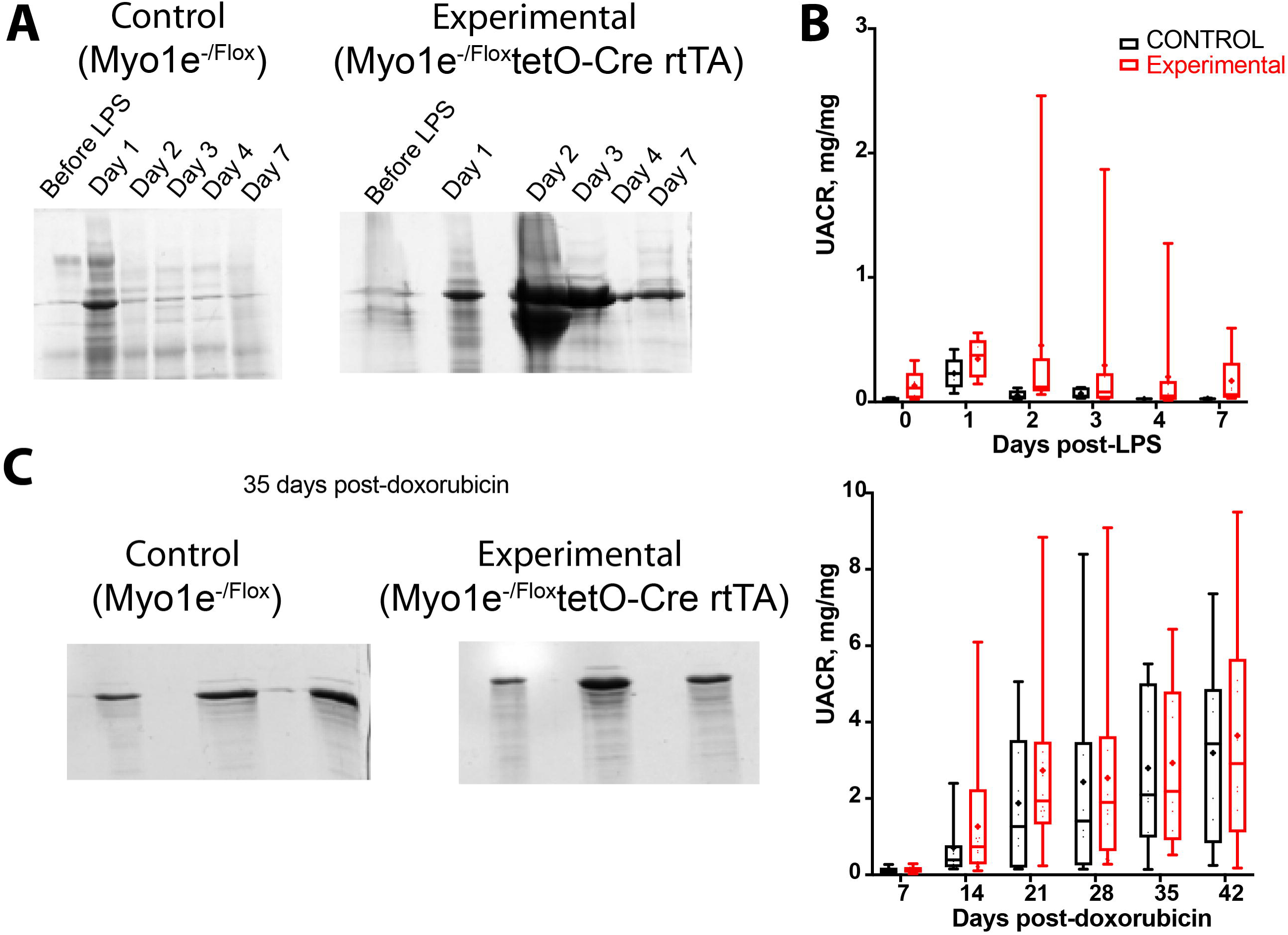
LPS-induced transient glomerular injury and doxorubicin-induced nephropathy in Myo1e-KO mice leads to slightly elevated proteinuric response compared to control mice. A,B. Control and Myo1e-KO mice were treated with doxycycline for 2 weeks starting at the age of 8 weeks. 3 weeks after completion of doxycycline treatment, mice were subjected to LPS treatment to induce proteinuria and followed for 7 days. Urine samples from one control and one KO mouse collected throughout the 7-day period were separated by SDS-PAGE and stained with Coomassie (A). Urinary albumin/creatinine ratio measured in urine samples collected from multiple animals (5 control and 9 KO) at the indicated time points in plotted in B. C,D. Control and Myo1e-KO mice were treated with doxycycline for 2 weeks starting at the age of 8 weeks. 3.5 weeks after completion of doxycycline treatment, mice were subjected to doxorubicin treatment. The Coomassie-stained SDS-PAGE gel in C shows proteins present in urine samples collected from 3 control and 3 KO mice. Quantification of the urinary albumin/creatinine ratio is shown in D (12 control and 15 KO animals). Boxes in B and D extend from the 25th to 75th percentiles, the line in the middle of each box corresponds to the median, and the whiskers extend down to the smallest value and up to the largest. All data points are shown.

As a second model of podocyte injury, we used the Adriamycin nephropathy model, which causes loss of podocytes (Hakroush et al., 2014). For these experiments, we used an intermediate dose of Adriamycin. Adriamycin injection caused a gradual increase in proteinuria, which was comparable in both control and experimental mice. Long-term follow up showed no pronounced histological changes in the experimental mice compared to control mice (data not shown).

## Discussion

In the current paper, we set out to determine whether inactivation of myo1e in adult podocytes can cause a significant disruption of the renal filtration barrier or reduce the ability of the glomerular filter to recover from an injury. This question was prompted by several considerations. First, with a number of studies showing that myo1e may be involved in assembly and modulation of cell-cell and cell-substrate contacts (Bi et al., 2013; Heim et al., 2017), podocytes lacking myo1e may be unable to maintain their foot process architecture, resulting in foot process effacement and proteinuria. Second, since podocytes have the ability to adjust to the changing capillary size or reduced podocyte number by altering their shape and area, cytoskeletal proteins, such as myosin 1e, could play an important role in this compensatory process. Furthermore, recovery from podocyte injury and reestablishment of foot processes and slit diaphragm complexes may rely on podocyte motility, which, in turn, may depend on myosin activity. Finally, since our previous study has shown that myo1e contributes to breast cancer progression (Ouderkirk and Krendel, 2014), inhibition of myo1e may be an attractive therapeutic approach for cancer treatment, therefore, it is important to establish whether the loss of myo1e activity may have serious adverse effects on kidney function.

Using an inducible knockout system, we have found that knockout of myo1e in both developing and adult podocytes causes proteinuria. The magnitude of proteinuria upon perinatal knockout of myo1e was similar to that in our earlier study that used a non-inducible Cre transgene (NPHS2-Cre) to cause myo1e knockout in developing podocytes (Chase et al., 2012). Comparing the extent of proteinuria in the current study in the adult vs. perinatal myo1e knockout shows that knockout of myo1e in the adult kidney leads to lower levels of proteinuria. This observation is in line with our previous finding that the level of urinary albumin excretion is much higher in mice with the germ-line knockout of myo1e compared to those with podocyte-targeted knockout. As we have previously hypothesized, at least one of the possible explanations for that observation is that myo1e expression in podocytes may be especially important during early stages of glomerular development, before NPHS2 promoter-driven Cre expression occurs. This hypothesis is confirmed by our observation that proteinuria was lower when the myo1e knockout performed in adult mice. Overall, the loss of myo1e from adult podocytes appears insufficient to cause nephrotic-range proteinuria or glomerulosclerosis. This observation provides a favorable basis for attempting to use myo1e inhibition as a therapeutic approach in cancer treatment.

We then asked whether the lack of myo1e impairs the ability of podocytes to recover from injury. We used two models of glomerular injury, low-dose LPS injection, which damages podocytes via actin reorganization triggered by the activation of Toll-like receptor 4 (Reiser et al., 2004) and Adriamycin injection. We hypothesized that if myo1e is necessary to maintain normal foot process organization in adult podocytes, its knockout may exacerbate podocyte injury induced by LPS. On the other hand, LPS treatment increases podocyte motility (Wei et al., 2008), thus, it may lead to foot process effacement via activation of the cytoskeletal pathways that normally are involved in cell migration or spreading. If that is the case, myo1e knockout could also be protective against LPS-induced podocyte motility. We observed a slight increase in LPS-induced proteinuria and no dramatic difference in recovery from this transient injury in mice with the induced knockout of myo1e, indicating that myo1e knockout did not have a protective effect on foot process effacement and did not significantly impair recovery. Similar results were obtained using a different model of glomerular injury, Adriamycin injection.

Overall, our study shows that myo1e removal from adult podocytes causes disruption of selective glomerular filtration and foot process effacement. However, the effects of myo1e loss from the adult kidneys are less dramatic than the absence of myo1e during renal development.

## Acknowledgements

This work was supported by the National Institute of Diabetes and Digestive and Kidney Diseases Award R01DK083345 to M.K. The content is solely the responsibility of the authors and does not necessarily represent the official views of the National Institutes of Health.

