## Supplemental Figure 1 for "Myo1e knockout in the adult podocytes leads to proteinuria but has less severe consequences for kidney function than Myo1e loss during renal development"

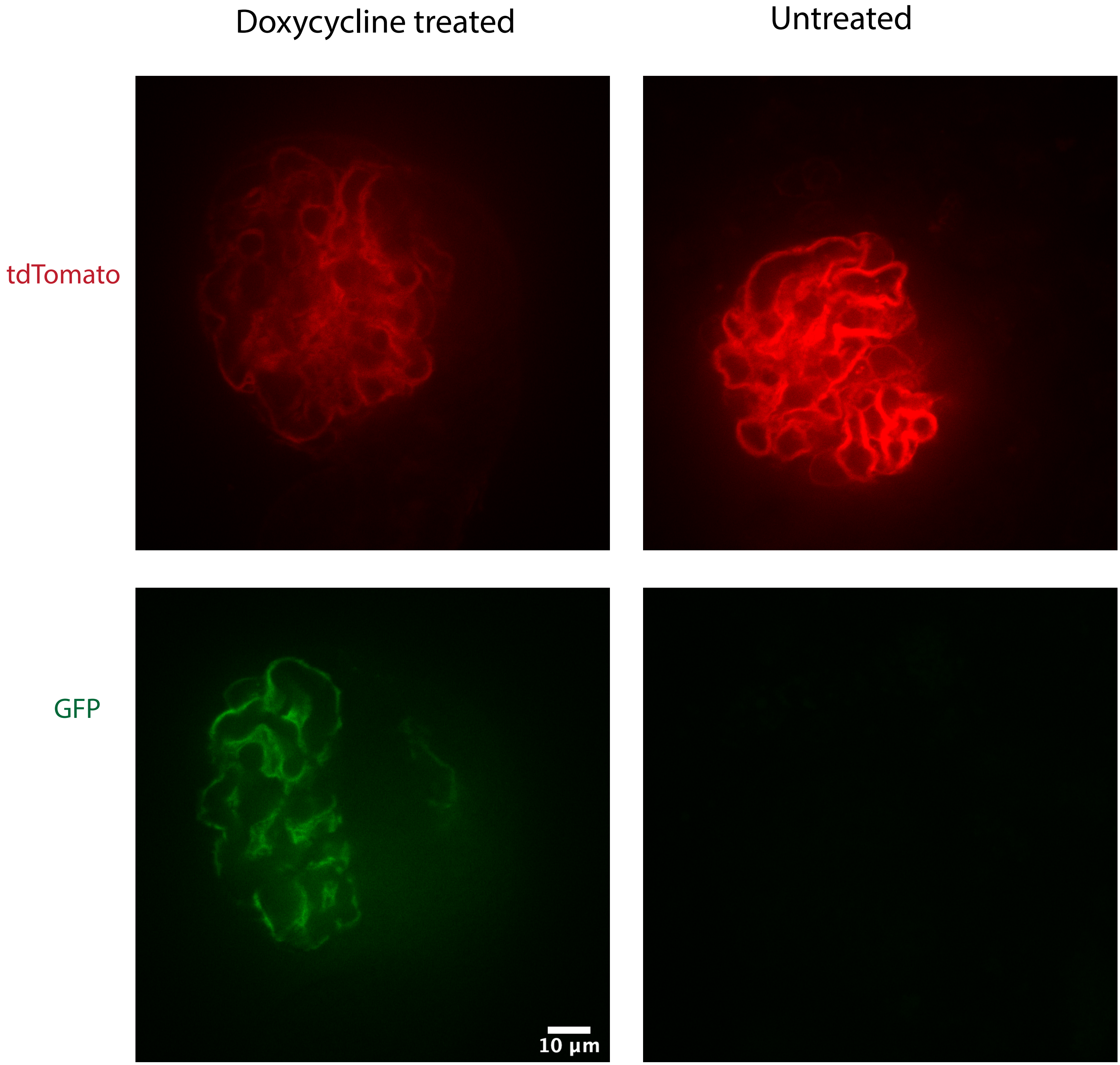


Figure S1. Glomeruli isolated from tetO-cre NPHS2-rtTA2 mTmG mice treated with doxycycline at the age of 8 weeks exhibit expression of GFP in podocytes. Glomeruli isolated from mice not treated with doxycycline express only tdTomato.
