## Supplemental Figure 2 for "Myo1e knockout in the adult podocytes leads to proteinuria but has less severe consequences for kidney function than Myo1e loss during renal development"

**A**

Control (Myo1e$^{-/Flox}$) Experimental (Myo1e$^{-/Flox}$tetO-Cre rtTA)

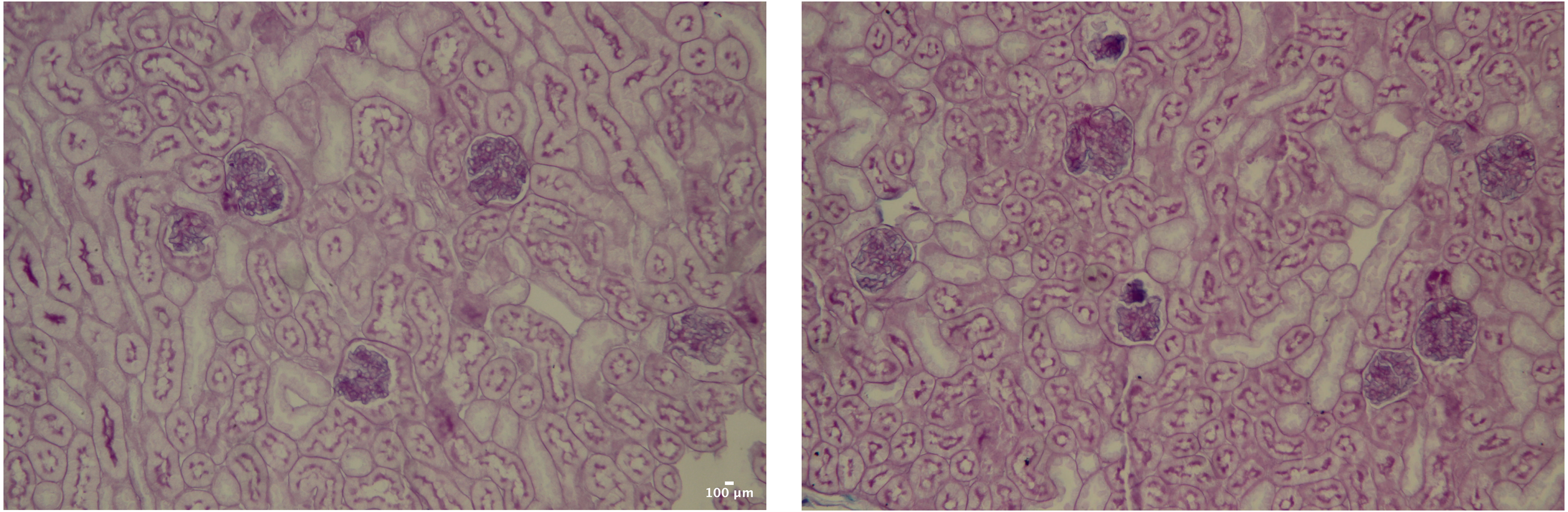


Figure S2. PAS staining of kidney sections obtained from doxycycline-treated control and Myo1e-KO mice. Doxycycline treatment was performed at the age of 8 weeks.
